# HiTea: a computational pipeline to identify non-reference transposable element insertions in Hi-C data

**DOI:** 10.1101/2020.04.27.060145

**Authors:** Dhawal Jain, Chong Chu, Burak Han Alver, Soohyun Lee, Eunjung Alice Lee, Peter J. Park

## Abstract

Hi-C is a common technique for assessing three-dimensional chromatin conformation. Recent studies have shown that long-range interaction information in Hi-C data can be used to generate chromosome-length genome assemblies and identify large-scale structural variations. Here, we demonstrate the use of Hi-C data in detecting mobile transposable element (TE) insertions genome-wide. Our pipeline HiTea (**Hi**-C based **T**ransposable **e**lement **a**nalyzer) capitalizes on clipped Hi-C reads and is aided by a high proportion of discordant read pairs in Hi-C data to detect insertions of three major families of active human TEs. Despite the uneven genome coverage in Hi-C data, HiTea is competitive with the existing callers based on whole genome sequencing (WGS) data and can supplement the WGS-based characterization of the TE insertion landscape. We employ the pipeline to identify TE insertions from human cell-line Hi-C samples. HiTea is available at https://github.com/parklab/HiTea and as a Docker image.

## INTRODUCTION

Over half of the human genome is composed of repetitive DNA sequences(de Koning *et al*., 2011). The repeats belong to two major classes: *(i)* tandem repeats, consisting of DNA sequences from few bases to few hundreds of bases that have expanded in tandem, stretching up to millions of bases in the genome; and *(ii)* transposable elements (TEs), interspersed throughout the genome and accounting for 44% of the human genome(Mills *et al*., 2007). Unlike tandem repeats, TEs are capable of transposition, in which they move from one genomic location to another. The distinct self- or *trans-* encoded mechanisms used by the TEs for transposition are used to group them into several families(Wicker *et al*., 2008). Although a vast majority of the TEs are inactive, a small fraction (<0.05%) still remains active in the human genome(Mills *et al*., 2007), primarily SINEs (Small Interspersed Nuclear Elements), LINEs (Long Interspersed Nuclear Elements), and SVAs (SINE-VNTR-*Alu*).

The transposition events are a major source of genomic structural variation (SV) and play an important role in a multitude of human genetic diseases(Hancks and Kazazian, 2016). For example, elevated levels of non-reference L1Hs (LINE) insertions are associated with epithelial carcinomas(Hancks and Kazazian, 2016; Lee *et al*., 2012; Chenais, 2015); Alu (SINE) insertions are associated with cystic fibrosis and hemophilia(Chen *et al*., 2008; Vidaud *et al*., 1993); and a recent case of Batten’s disease that led to the development of an individualized antisense oligonucleotide therapy(Kim *et al*., 2019) was caused by an SVA insertion. The TE sequences may also encode a range of regulatory features such as promoters, enhancers, transcription factor binding sites, and non-coding regulatory RNA transcripts(Chuong *et al*., 2017). Thus at the molecular level, transposition can result in altered gene expression, splicing/RNA stability defects, genome instability, or decreased integrity of centromere and telomeres(Bourque *et al*., 2018).

In particular, TE sequences are a rich source of binding sites for an insulator protein CTCF, which plays a key role in regulating the 3D structure of chromatin. The extended loops of the DNA are maintained by binding of CTCF at the base of the loop; indeed, the Hi-C chromatin maps suggest enrichment of SINE elements at the topologically associated domains (TAD) boundaries(Rao *et al*., 2014). The TE-derived CTCF binding sites are a fundamental source for mammalian genome evolution at various time scales, with some highly conserved across species and some species-specific expansions of CTCF sites co-occurring with species-specific TADs(Schmidt *et al*., 2012; Cournac *et al*., 2016). Given the important regulatory role of TEs(Ayarpadikannan and Kim, 2014; Garcia-Perez *et al*., 2016; Ahmed and Liang, 2012), identification of their transposition is important in understanding the disease biology, gene regulation, and 3D chromatin organization.

Several computational tools are available for identifying non-reference (either somatic and germline) TE insertions from WGS data(Rishishwar *et al*., 2017). A key component of such methods is the identification of discordant read pairs (RP), whose genome alignments display unexpected between-pair distance or orientation. A discordant RP with one end mapping to the consensus TE sequence and the other end mapping to the reference genome is indicative of a TE insertion. Discordant RPs are typically accompanied by ‘clipped’ reads, whose partial alignment can be used to obtain base-pair resolution of the breakpoints. With judicious integration of these criteria and appropriate thresholds, candidate TEs insertions can be predicted across genome.

Besides WGS, another data type that involves a large amount of sequencing is Hi-C, an unbiased genome-wide extension of the chromosome conformation capture technique. Hi-C experiments(Rao *et al*., 2014; Schmitt *et al*., 2016) are conducted primarily to understand the long-distance regulatory relationships in the genome (e.g., which enhancer interacts with which promoter). In this experiment, the cross-linked DNA fragments are first digested with a suitable restriction endonuclease (RE). Then, random ligation is performed in a condition that favors ligation between cross-linked fragments. The resulting ligation product contains pairs of fragments that were close in 3D proximity. Sequenced Hi-C reads indeed show that the effective insert sizes—the distance between the mapped mates—range from few hundred to millions of bases. Consequently, the proportion of discordant RPs, that are <20% in WGS, are in the excess of 50-70% for Hi-C data. Furthermore, as the sequenced fragments are generated post-ligation step, the proportion of reads carrying split mapping (due to encompassed RE sites) is higher in the Hi-C data. These features thus limit the use of WGS-based TE detection tools on Hi-C data.

Here, we present a computational pipeline HiTea (**Hi**-C based **T**ransposable **e**lement **a**nalyzer), which identifies non-reference TE insertions of the LINE, SINE and SVA families using Hi-C data. Our comparisons show that HiTea (run on Hi-C) performs similarly to a commonly-used WGS-based tool (run on WGS at similar coverage)(Gardner *et al*., 2017). With increasing realization of 3D chromosomal structure as a regulatory component of gene regulation, large scale efforts such as 4D Nucleome(Dekker *et al*., 2017) are underway to aim to map genome organization across cell-types and disease models. Our results indicate that Hi-C data can be used not only to study 3D genome organization but also to characterize the non-reference TE insertions.

## METHODS

### Informative Hi-C read pairs for non-reference TE detection

To understand the methodology underlying HiTea, we first describe the different types of read pair (RP) mappings observed in Hi-C data (Fig.1A). Discordant RPs, defined in paired-end sequencing, are RPs with unexpected distance or orientations between paired mate reads when mapped to the reference genome. Due to the intrinsic design of Hi-C experiments for detecting interactions between two distant genomic loci, a major proportion of RPs (typically 50-70%) in Hi-C data are discordant with large (>1kb) mapping distances or atypical orientations of the paired mates. A small proportion (6-30%) of RPs display WGS-like concordant read mapping configuration (Fig. 1A, panel i), where both mates map close (< 500bp) to each other in convergent orientation.

**Figure 1:**
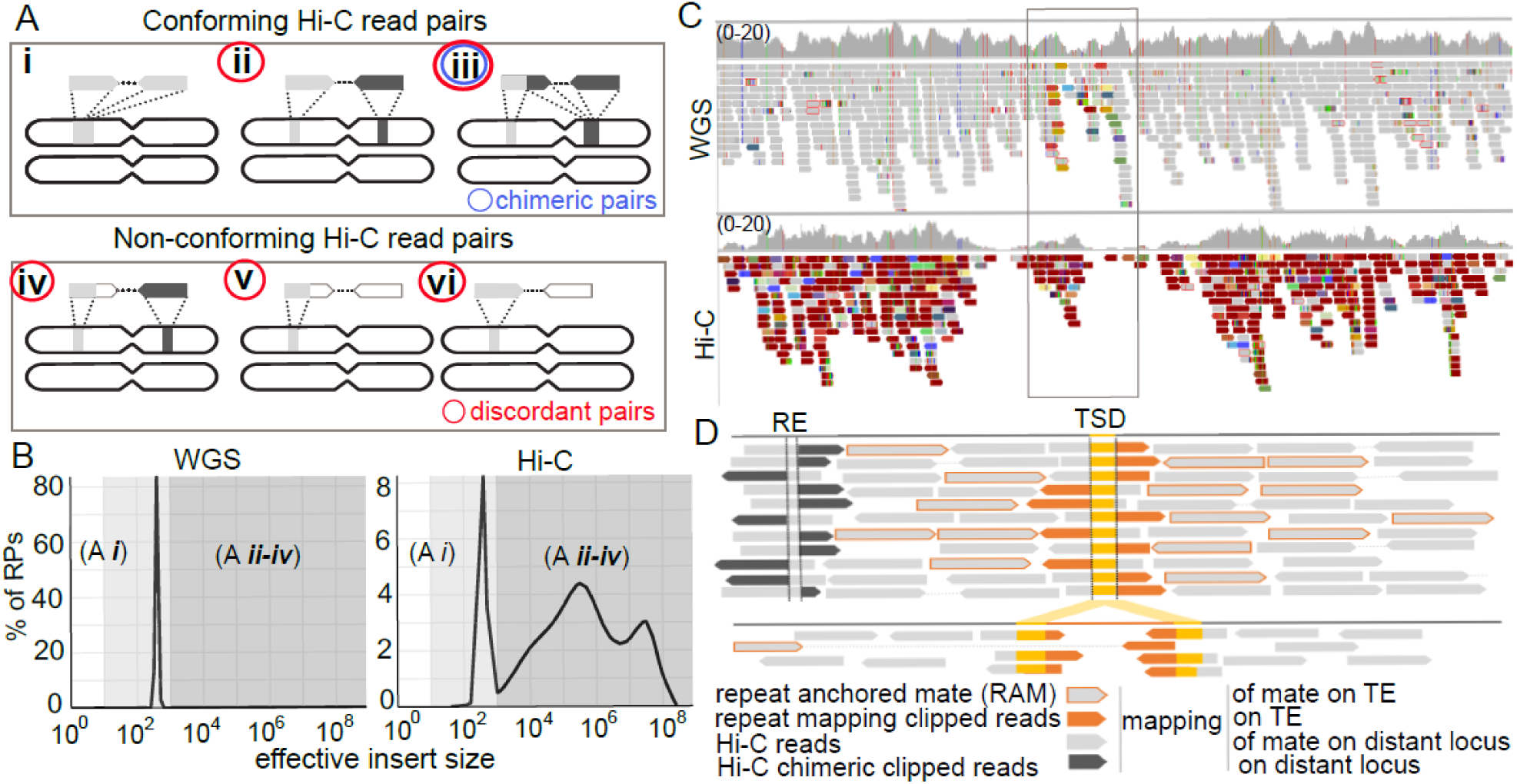
Properties of Hi-C reads supporting a TE insertion. **(A)** Hi-C read pairs (RPs) can be grouped into two classes that we termed ‘conforming’ and ‘non-conforming’. Conforming RPs comprise of *(i)* WGS-like pairs with short insert sizes, *(ii)* pairs with large effective insert sizes, and *(iii)* chimeric RPs where the clip-sequence maps convergent to mapped locus of its paired mate. Non-conforming RPs comprise of mapping configurations where *(iv)* the clipped sequence does not display chimeric mapping or *(v-vi)* the mate remains unmapped on reference genome. **(B)** Comparison of the between-pair distances for WGS and Hi-C experiments. **(C)** A genome browser view of a true insertion event, showing both coverage and the discordant RPs (non-gray color) in WGS and Hi-C experiments. Box marks the TE-insertion site. Mapped read pairs in the display are color-coded by the insert sizes using default IGV color scheme. **(D)** A schematic of Hi-C read configuration at insertion site. Clipped reads supporting TE insertion exhibit partial mapping to TE-family consensus (orange), whereas those that do not, map at distant reference locus (black). RPs with a mate mapping to the TE-family consensus are displayed with orange outline. (RE: restriction endonuclease, TSD: target site duplication)

The RPs in Hi-C data can also be classified into two different categories. First, we introduce the terminology *conforming* RPs to refer to those with mapping configuration explained solely by the Hi-C experiment. For instance, conforming RPs with unique mapping of the entire mate reads on two proximal or distant genomic loci are prevalent in Hi-C data (Fig. 1A-i,ii). Here, the between-pair distance can range from WGS-like insert size (*i.e*., ∼500bp) to millions of bases (Fig. 1B). A third type of conforming RPs are those in which the 5’ portion of a mate maps uniquely to the genome and the 3’ portion maps convergent on the genomic locus of the matching mate, and the two portions are connected with the RE ligation motif (Fig.1A-iii). These mappings are referred to as chimeric Hi-C pairs (∼10-20%) and are included in the 3D-contact matrices. Second, the remaining RPs (∼10-30%) do not conform to any expected configuration of read mappings, and thus are discarded in standard analyses. In those *non-conforming* RPs, one or both mates remain unmapped, multi-mapped, or their partial mapping does not produce chimeric Hi-C pairs (Fig. 1A-iv,v,vi). To identify non-reference TE insertions, HiTea uses non-conforming RPs whose partial (clipped) sequences or one entire mate read map to TE sequence assemblies.

In Fig. 1C, we show the distribution of reads along a small genomic region. In WGS data, the genomic coverage is relatively even. In Hi-C data, the coverage is more variable; however, much of the region is still covered with at least some reads, thus allowing for the possibility that most TE insertions can be captured. The proportion of discordant RPs (non-gray colors) is very high in Hi-C data.

### Identification of TE insertion breakpoints

HiTea starts by identifying non-conforming RPs using Pairtools (https://github.com/mirnylab/pairtools). In the discovery step, the clipped reads without legitimate RE-ligation motif are then mapped (using BWA-MEM(Li and Durbin, 2010) with ‘-a -k 13 -T 20’) to family-wise TE consensus assemblies published earlier(Gardner *et al*., 2017) for Alu (SINE), L1Hs (LINE) and SVA (https://melt.igs.umaryland.edu/downloads.php). Additionally, it uses a separate 200 base long PolyA sequence to improve detection sensitivity of TEs, especially those with long PolyA tails. For the alignment, we note that many polymorphic insertions may have sequences distinct from the family-based consensus. To accommodate such cases, HiTea offers an option to remap clipped reads that initially fail to map to a TE family consensus, to a user-provided set of polymorphic sequences for a TE-family or sequences of the members of its subfamily (e.g., from Repbase(Bao *et al*., 2015)). HiTea, in principle, can also detect insertions of other template-based transposons such as an active human endogenous retrovirus (HERV-K), as long as adequate TE-consensus sequences are provided.

The clipped sequences are derived from non-conforming Hi-C RPs, where minimum clip length (default: -s 20) can be defined by the users. Using a two base-pair leeway, a breakpoint on the reference genome is determined as the location with the maximum number of clipped reads at a locus (Supl.Fig.1). HiTea simultaneously records all non-conforming RPs in which a read maps to the reference genome and its ‘anchor’ mate maps to the TE-consensus assembly (using default BWA-MEM settings). We refer to these as **R**epeat-**A**nchored non-conforming Hi-C **M**ates (RAMs) pairs (Fig. 1D), following the terminology introduced earlier(Lee *et al*., 2012). All breakpoints supported by at least two clipped reads with partial mapping to a TE-consensus are further interrogated for enrichment of available TE supporting clipped reads and RAMs using a negative binomial model (Supl.Fig.1). The candidate sites where the numbers of clipped reads and RAMs are less than 5% and 2.5%, respectively, of the total Hi-C coverage at the locus are omitted as unreliable.

Unlike WGS, where the RAM pairs are clustered around the sites of TE-insertion, Hi-C data exhibits wider mapping area. Though, both WGS and Hi-C data are biased by GC-content or overall mappability, the coverage in Hi-C is additionally clearly biased by the density of RE sites at the locus. Hence, HiTea uses a negative binomial model to assess the enrichment of TE-insertion supporting reads (i.e., RAM pairs and clipped-reads) at the locus. To model the biases, HiTea uses randomly selected loci in the genome that have similar coverage of the non-conforming RPs as the site under investigation. Then, the count of TE-supporting reads at a locus is assessed against negative binomial model built from the random set.

### Filtering and annotation of non-reference TE insertions

A substantial fraction of clipped reads in Hi-C data display chimeric mapping (Fig. 1A, panel iii) carrying a ligation motif at the clip position. To avoid calling such canonical Hi-C interactions as TE insertions, HiTea filters out insertion candidates whose predicted breakpoints on either the reference genome or TE-consensus are within 3-bases (user-defined) of the ligation motif (Fig.1D, clip reads at RE site; Supl.Fig.1 for detailed filtering steps). It also filters out candidates when multiple breakpoints are predicted around a putative breakpoint, as it is likely to be a complex variant other than a TE insertion. At the sites of insertion, clipped mapping positions of the reads indicate a breakpoint where reads mapping to the reference genome cluster (Fig. 1D). HiTea expects that the genuine breakpoint should also show reciprocal cluster of the clipped sequences when mapped to the TE-consensus. Insertions defying this expectation are removed as invalid. Furthermore, insertions where clipped reads mapping only to the PolyA sequences are omitted as potential simple repeat expansions. The genuine breakpoints are expected to have clip-sequences mapping to PolyA sequence or presence of a degenerate polyA sequence (here we look for a stretch of 7 As or Ts in the proximal 10 bases at the breakpoint on clipped sequences). Subfamily annotation of the insertion is done by mapping the longest clipped sequence to the subfamily consensus sequence derived from Repbase(Bao *et al*., 2015). HiTea further detects target site duplication, strand information, and estimates the size of insertion from the observed mapping of the clipped sequences on the TE-consensus. HiTea is written in PERL and R. It uses GNU-parallel(Tange, 2011) for parallelization over available cores. The insertions are reported in bed format, with following status. Status-3 insertions are supported by right- and left-hand side mapping of the clipped reads (Fig. 1D), whereas status-2 insertions represent a subset of status-3 cases that overlap the reference copy of the same TE family. If the insertion is supported by clipped reads at one side but have unmapped reads on the other site with polyA stretches (as defined earlier), such instances are flagged with status-1.

## RESULTS

### HiTea shows performance comparable to that of a WGS-based method

To assess the performance of HiTea, we utilized Hi-C data generated from the HapMap cell line GM12878(Rao *et al*., 2014). This cell line has been extensively characterized using a wide range of technologies and sequencing platforms. To generate the gold standard for comparison, we used an improved version of our algorithm Tea(Lee *et al*., 2012) on PacBio HiFi long reads(Zook *et al*., 2016) with extensive manual curation (hereafter referred to as the PacBio reference). For WGS, we employed Mobile Element Locator Tool (MELT)(Gardner *et al*., 2017), a popular software package with reportedly superior performance at moderate sequencing depth(Rishishwar *et al*., 2017). The full datasets consisted of ∼5B RPs for Hi-C(Rao *et al*., 2014) (MboI-digested dataset; downloaded from 4DN data portal) and ∼1.4B RPs for WGS (downloaded from the 1000 Genomes project). Sequencing depths have considerable impact on the precision and recall(Rishishwar *et al*., 2017), thus we randomly down-sampled Hi-C data to1.4B RPs (∼80X coverage) to provide a fair comparison between platforms. At this coverage, 79% of the genome in WGS and 57% in Hi-C data are covered with at least 60X coverage (Fig. 2A). The coverage was calculated by counting reads with mapping quality of at least 10 (MAPQ ≥10).

**Figure 2:**
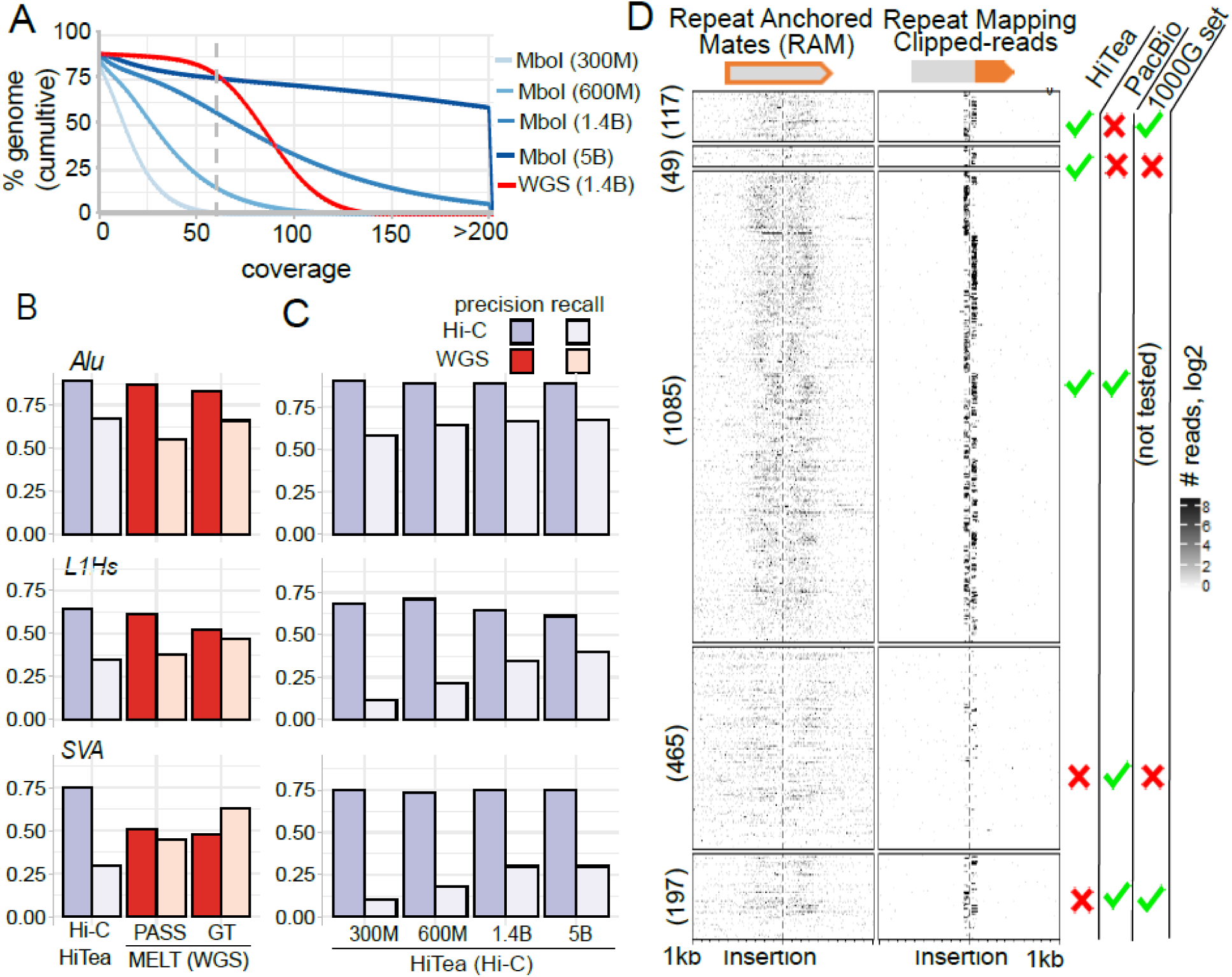
Performance of HiTea. **(A)** Cumulative distribution of the coverage for different datasets. Gray dotted line marks 60X coverage. **(B)** Precision and recall for detecting insertions of Alu, L1Hs and SVA families using HiTea (on Hi-C) and MELT (on WGS) at 1.4B sequencing depth. PASS and GT refer to the more and less stringent call sets, respectively, in MELT. **(C)** Precision and recall comparison at different sequencing depths of Hi-C experiment. **(D)** 5’ end coverage for the RAMs whose mates map to the TE consensus (left) or reads whose clipped-sequences map to the TE consensus (right). The insertions are grouped according to the criteria shown on the right. PacBio is the reference set constructed using PacBio HiFi reads; 1000G set refers to insertions detected in the 1000 Genome data by MELT.

The candidate insertions predicted by HiTea (ran on Hi-C data) and MELT (ran on WGS data) were compared against the PacBio reference set (Fig. 2B). We used two sets of insertions reported by MELT for GM12878: *(i)* the stringent “PASS” set (1122 insertions, referred as MELT-PASS) and *(ii)* a more lenient set that includes the PASS variants and others for which genotype could still be inferred (1443 insertions, referred as MELT-GT) in the comparisons. A total of 1251 insertion were identified by HiTea while the PacBio reference set consisted of 1747 insertions.

Overall, HiTea correctly identified 1085 insertions (Fig 2B). The precision (fraction of the true positives among all identified insertions) was 0.87; recall (fraction of true positives among all positives) was 0.62 with F1 score of 0.72. MELT-PASS and MELT-GT correctly recovered 925 (precision 0.82, recall 0.53, F1 0.64) and 1115 (precision 0.77, recall 0.64, F1 0.7) insertions, respectively.

i. *Alu*. Most of the insertions were Alu, as expected. Among the 1493 Alu insertions from our reference set, HiTea correctly identified 1000 (precision 0.89, recall 0.67, F1 0.76) insertions from the Hi-C data. Whereas, MELT-PASS correctly identified 825 (precision 0.87, recall 0.55, F1 0.68) and MELT-GT recovered 986 (precision 0.83, recall 0.66, F1 0.74) Alu insertions from the WGS data. These results suggest that HiTea (ran on Hi-C) has considerably better performance at detecting Alu compared to MELT (ran on WGS) (Fig. 2B). Notably, HiTea can detect Alu insertions with competitive precision and recall from Hi-C samples with lower coverages (Fig. 2C). For instance, at 600M RPs (∼40X sample; recommended sequencing depth by the 4DN consortium) and 300M RPs (∼20X coverage), the precisions are nearly uniform (i.e. 0.89 for 1.4B, 0.89 for 600M and 0.90 for 300M) and the recalls decrease only slightly, from 0.67 (1.4B, F1 0.76) to 0.65 (600M, F1 0.75) and 0.59 (300M, F1 0.71) (Fig. 2C). We compared the proportions of the clipped reads, which are the starting point of TE insertion identification in HiTea, and RAM reads that map to Alu consensus between Hi-C (identified by HiTea) and WGS (identified by MELT) at the equal sequencing depth of 1.4B. Although the proportions of Alu-mapping clipped reads (44% in Hi-C and 53% in WGS) were higher, we observed that the proportion of RAMs pairs mapping to the Alu consensus is much higher for Hi-C (43% of total RAMs) than WGS (13% of total RAMs). Taken together, better proportions of mapping of clipped and RAM reads in Hi-C is likely associated with better performance of HiTea on Alu.
ii. *L1Hs*. Our PacBio reference set contained 194 high-confidence L1Hs insertions. HiTea correctly identified 67 (precision 0.64, recall 0.35, F1 0.45), whereas MELT-PASS and MELT-GT detected 73 (precision 0.61, recall 0.38, F1 0.47) and 91 (precision 0.52, recall 0.47, F1 0.49), respectively (Fig. 2B). With respect to sequencing depths, recall increased as the depth increased, from 0.11 for 300M (F1 0.19) to 0.4 for 5B RPs (F1 0.48), while the precision remained in a similar range (0.61 to 0.71) (Fig.2C). Interestingly, the proportions of both clipped and RAM reads mapping to the L1Hs consensus were substantially higher in the WGS data (39.5% and 84.2% respectively) compared to the Hi-C data (27.5% and 52.7% respectively). Transposed copies of L1Hs are frequently associated with 5’ truncation and/or inversion. Moreover, during target-primed reverse transcription (TPRT), L1 RNA often accommodates sequences from the downstream genomic region(Pickeral *et al*., 2000). These additional features may lower the performance of HiTea for L1Hs compared to Alu.
iii. *SVAs*. HiTea has relatively poor sensitivity towards SVAs. Of 60 SVAs in the PacBio reference set, HiTea correctly identified 18 (precision 0.75, recall 0.3, F1 0.43), whereas MELT-PASS and MELT-GT respectively detected 27 (precision 0.51, recall 0.45, F1 0.48) and 38 (precision 0.48, recall 0.63, F1 0.55) instances. Although the proportions of RAMs mapping on the SVA-consensus were comparable (2.7% for Hi-C vs 2.5% for WGS), the proportions of SVA mapping clipped reads were substantially different (4.6% in Hi-C vs 7.1% in WGS). SVAs comprise of frequently expanded hexameric repeats at the 5’, variable number of tandem repeats (VNTR) in the middle, and Alu-like sequences at the 3’. This complex structure may lead to the relatively poor mapping of SVA-originating reads to the SVA consensus (*e.g*., some SVA reads map to the Alu consensus instead), and thus affect the performance of HiTea for SVAs. Nonetheless, the precision of detecting SVAs was strikingly high for HiTea (0.73 to 0.75) as compared to the MELT calls (<0.51) (Fig. 2B,C). The impact of sequencing depth for SVAs was similar to that for L1Hs.

Of the 1251 HiTea insertions (at 1.4B), ∼13% (166) did not overlap with the PacBio reference set. Hence, we interrogated them against a collection of 1000 Genome TE insertion set (at a population allele frequency ≥ 10%; results were similar for AF ≥0.01% and AF ≥0.1%), identified on the low coverage WGS data by MELT(Gardner *et al*., 2017). Our comparison suggested that 117/166 (∼71%) HiTea-specific insertions overlap with the population-based TE-insertion set, suggesting that these are true insertions missed by the PacBio reference set. This also suggests that the precision and recall measures above represent lower bounds.

HiTea missed ∼38% (662/1747) of the insertions from the PacBio reference set. Of the 662, 197 insertions overlapped with 1000G set. We assessed the 5’ end coverage of RAMs whose mates or clipped sequences map to the TE consensus in Fig. 2D. This coverage plot(Gu *et al*., 2018) shows that the missed events by HiTea do not have a sufficient number of clipped reads (lower right panels in Fig. 2D; 381/465 and 95/197 have less than two non-Hi-C chimeric clipped reads mapping to the TE-consensus at the locus.

Since the 1.4B-RPs datasets used above are larger than typical datasets, we repeated the above analysis with down-sampled datasets with ∼600M RPs (∼35-40X). Our comparison suggests that HiTea (run on Hi-C) shows consistently higher precision in detecting Alu, SVA and L1Hs compared to MELT (run on WGS data) (Supl.Fig.2A). A total of 1016/1152 HiTea insertions (precision 0.88, recall 0.58, F1 0.70) and 908/1134 MELT-PASS insertions (precision 0.80, recall 0.52, F1 0.63) overlapped with the PacBio reference set (Supl.Fig.2A). The insertions missed by HiTea did not seem to show clip-read coverage at the respective loci (Supl.Fig.2B).

We tested HiTea on a range of human Hi-C datasets generated using different REs. A 4-cutter RE (MboI, DpnII) is expected to cut the DNA at every 256 bases whereas a 6-cutter (HindIII, NcoI) will digest the DNA at 4096bp on average. The infrequent cuts by a 6-cutter are expected to provide low spatial resolution of the Hi-C (Supl.Fig.3A), resulting in a smaller number of clipped reads along the genome. Indeed, when Hi-C datasets generated using different REs for GM12878 cell line(Rao *et al*., 2014) were compared, the overall recall dropped from 0.62 (MboI digested Hi-C, 1.4B RPs, F-score 0.72) to 0.41 (1.8B RPs, HindIII digested Hi-C, F-score 0.56). For comparison, the overall recalls for WGS sample were 0.53 (MELT-PASS, F-score 0.64) and 0.64 (MELT-GT, F-score 0.7) at 1.4B RPs (Supl.Fig.3B). Nonetheless, HiTea showed a high precision (0.88) compared to MELT-PASS (0.82) and MELT-GT (0.77). Besides 17 unique, remaining 794 (98%) insertions from the HiTea run either overlapped with PacBio reference set or the 1000G set, whereas about 79% (811/1032) of the missed insertions displayed poor coverage of clipped reads (Supl.Fig.3C, D). With the decreasing sequencing cost, many studies are now using either a 4-cutter or a mix of 4-cutter enzymes, and these high-resolution Hi-C datasets will be suitable for HiTea analysis.

Next, we assessed the performance of HiTea on another widely-characterized cell line, K562. We obtained WGS and Hi-C (MboI digested Hi-C) data from Cancer Cell Line Encyclopedia project(Barretina *et al*., 2012) and a published study(Rao *et al*., 2014), respectively. As a PacBio reference set was not available for this cell line, we resorted to comparing the TE-insertions called by HiTea (on Hi-C) to those from MELT (on WGS). At comparable sequencing depth of 1.2B RPs between Hi-C and WGS data for K562 cells, a substantial fraction (769/958, ∼80%) of HiTea insertions overlapped with either MELT-derived (i.e. MELT-GT) insertions or 1000G set. In comparison, previously analyzed GM12878 (MboI digested, 1.4B RPs) exhibited similar (1101/1251, ∼88%) degree of overlap (Supl.Fig.4).

### HiTea aids in the characterization of the non-reference TE insertions

To assess whether HiTea can correctly identify insertions otherwise missed by MELT, we compared MELT-GT (better recall compared to MELT-PASS) and HiTea insertions (both at 1.4B RPs sequencing depth) using the PacBio reference set. Our analysis suggests that a substantial number of insertions overlapping with reference-genome copy of the same TE family are missed by MELT (Fig. 3A, B). TE detection along the reference TE copy of the same family can be challenging due to multiple reasons, such as poor mappability of the reads and structural variation within the reference-copies of the TE family. Therefore, several WGS-based tools filter out these insertions to limit the number of false positives(Ewing, 2015). However, when supporting reads are available and their mappings on both TE-consensus and reference genome provide sufficient confidence for the insertion, HiTea reports these events. Our reference set included 436 TE-insertions overlapping with the reference copies of the same TE-family. HiTea correctly identified 70 insertions reported in the PacBio reference, outperforming MELT (5 and 8 by MELT PASS and GT) (Fig. 3B).

**Figure 3:**
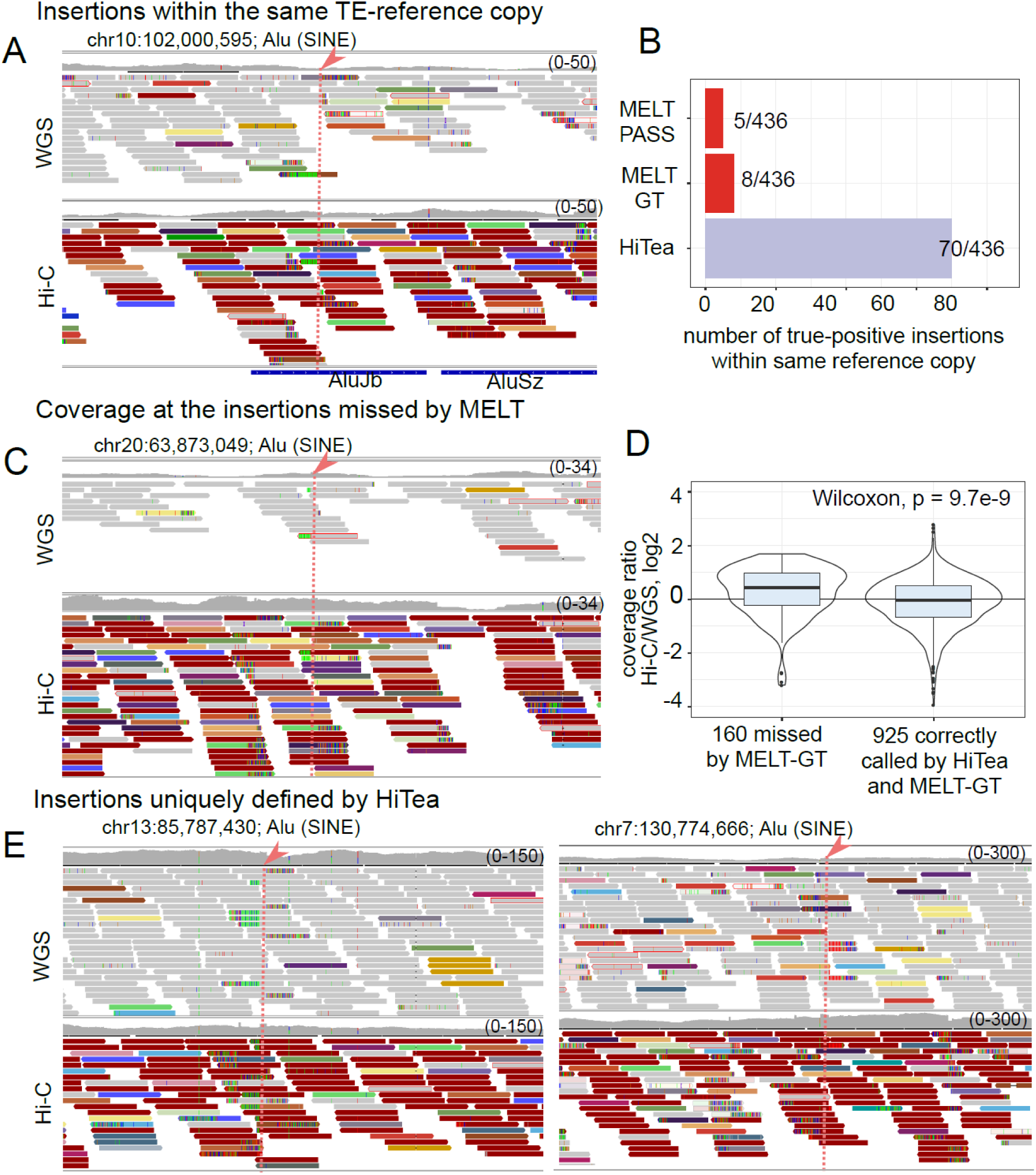
Examples of TE insertions detected in Hi-C but missed in WGS. **(A)** A browser view of an insertion overlapping the reference-genome copy of a TE-family. This insertion is identified by HiTea (on Hi-C) but missed by MELT (on WGS). Reads with concordant and discordant mapping configurations are displayed in gray and non-gray colors, respectively. The discordant RPs are color-coded according to their insert sizes. Dotted red line with arrowhead marks the insertion site. **(B)** Summary of TE insertions detected when the insertion occurs in the reference-genome copies of the TE-family. **(C)** Sequencing coverage comparison at an insertion correctly called by HiTea but missed by MELT. **(D)** The boxplot for the Hi-C/WGS read coverage ratios shows that Hi-C coverage is higher in cases identified by HiTea but missed by MELT-GT. **(E)** More examples of insertions called only by HiTea.

In total, HiTea identified 160 PacBio reference insertions missed by MELT-GT. Conversely, MELT-GT identified 180 insertions missed by HiTea from the reference set (Supl.Fig.5A, B). When assessed for the features that led to disqualification of these true-positive insertions by either MELT or HiTea, we observed that indeed insertions within a reference-genome copy of the same TE family were preferentially missed by MELT (66/160, ∼41%; Supl.Fig.5C). As the exact features used by MELT are unavailable (the code is not open source), we could not further investigate the instances missed by MELT. Over half of the insertions (124/180, ∼69%) missed by HiTea were due to poor coverage of clipped reads, proximity to the RE motif, coverage thresholds, and absence of clipped reads supporting polyA tails (Supl.Fig.5D).

Coverage in the Hi-C experiment is significantly higher around the RE sites in the genome. Thus, insertions proximal to the RE sites tend to have higher coverage of supporting reads even at relatively low overall sequencing depth. In the example shown in Fig.3C, the read coverage at an Alu insertion site missed by MELT-GT on chromosome 20 is much higher in Hi-C than in WGS, although the overall sequencing depth is the same (both bam files were subsampled to 10% of total reads for better visualization). To assess whether the same phenomenon is observed at many sites, we counted total 5’ end coverage in a 1kb window centered at the 925 insertions identified by both MELT-GT and HiTea and the 160 insertions identified only by HiTea. As expected, the insertions identified by both methods tend to have similar coverages, whereas those missed by MELT-GT tend to have relatively lower coverage overall in WGS compared to Hi-C (Fig.3D).

A total of 49/1251 (∼4%) insertions detected by HiTea were not explained by either the PacBio reference set or the 1000G set (Fig.2D, second panel from the top). Of these, 4 and 6 were reported by MELT-PASS and MELT-GT, respectively. These HiTea-specific insertions exhibit clear presence of TE-mapping clipped reads from Hi-C data (Fig.2D). Representative examples of two Alu insertions suggest that the HiTea-unique insertions have the support of both clipped and discordant reads at the insertion locus in the WGS data (Fig.3E). We suspect that many of these cases may be true positives that were missed by MELT due to its stringent filtering criteria.

### Installation and usage

HiTea is available at Github (https://github.com/parklab/HiTea) and as a Docker image (4dndcic/hitea:v1on Docker Hub). TE (Alu, L1Hs and SVA) family-wise consensus sequences and the genomic locations of the TE-family members required for running HiTea are provided for hg38 and hg19 human genome references, with a description on how to generate them for other types of TEs on the GitHub page. HiTea dependencies are PERL (≥ v5.24), R (≥ v3.2), bedtools (≥ v2.26)(Quinlan and Hall, 2010), samtools (≥v1.7), GNU-parallel(Tange, 2011) and Pairtools (https://github.com/mirnylab/pairtools). Additionally, there are mandatory (GenomicRanges, data.table, MASS) and optional (rmarkdown, knitr, EnrichedHeatmap(Gu *et al*., 2018), circlize) R packages used for computation and HTML-report generation steps respectively. Users can start the analysis with a single command by providing a name-sorted bam file, restriction enzyme used for the Hi-C assay and the genome build used to map the Hi-C data. HiTea auto detects if the read class information is present in the bam file (e.g. files obtained from 4DN data portal https://data.4dnucleome.org/ carry this information). If not, it automatically employs Pairtools to generate read class information. User-defined TE-consensus or polymorphic sequences and the genomic locations of the members of TE-sequences can be provided using a detailed input option. A HiTea run on a typical Hi-C dataset (∼600M RPs) takes about 3.5-4 hrs to complete with 8 cores and 20 G memory.

## DISCUSSION

Although used primarily for understanding three dimensional organization of the genome and its regulatory role, the long-range chromatin interaction information in Hi-C data have been used to assemble small scaffolds into chromosome-length assemblies(Dudchenko *et al*., 2017; Gong *et al*., 2018) and to identify copy number and translocations(Chakraborty and Ay, 2018; Dixon *et al*., 2018; Wang *et al*., 2020). In the present work, we have demonstrated that Hi-C can be used also to identify TE insertions.

The strong performance of HiTea was somewhat unexpected. Given the nature of the experiment, the read coverage for Hi-C is highly variable along the genome. We thus expected that there would not be enough reads at some TE insertions sites, resulting in degraded performance for HiTea compared to a WGS-based method. What makes HiTea competitive with a WGS-based method, however, is the use of clipped reads to locate candidate TE insertions at the discovery step, in contrast to the discordant RP-based candidate discovery in most WGS-based methods. The higher proportion of clipped reads (carrying no RE ligation junction) in Hi-C data (1.6%) than in WGS data (1.4%) is further helpful. Moreover, the proportion of RPs whose one end remains unmapped or multimapped is higher in the Hi-C data (21%) compared to the WGS data (14%) due to wider effective insert sizes, increasing the power of Hi-C data for detecting insertions. In particular, the TE insertions in the reference genome copies of the same family or those occurring in regions with comparatively lower coverage in WGS data are sometimes detected by HiTea but missed by MELT.

The availability of PacBio HiFi data (circular consensus sequencing method, with half the reads >50kb) for GM12878 made it easier to evaluate the performance of different methods. However, the TE insertion map based on this one sample is obviously incomplete, as seen by the fact that many HiTea candidates not present in the PacBio reference set were present in the 1000G data. A small fraction (<5%) of HiTea insertions were still not explained by either PacBio reference set or 1000G set. Although some of these insertion calls may be false positives, it is interesting to note that both WGS and Hi-C data show presence of discordant and non-conforming RPs mapping to the underlying TE consensus, respectively, along most of these loci. Additional long-read data or independent experimental validations may prove useful in discerning the nature of HiTea-specific calls.

The number of studies mapping chromatin organization in diverse organisms, cell types, and disease states as well as the collective efforts to organize such data have gained momentum(Dekker *et al*., 2017). However, it is imperative to mark structural variations in the genome before construing the chromatin interactions from Hi-C data as functional interactions, as we have demonstrated recently(Wang *et al*., 2020). HiTea exploits Hi-C data to identify non-reference TE insertions, using reads that otherwise would be discarded. Finally, although we compared call sets from Hi-C and WGS data in our analysis, the ideal scenario is to have both data types for a sample of interest, so that the insertions calls can be cross-validated and expanded. Continued development of more comprehensive reference TE insertions maps and robust computational methods for TE identification will be important.

## Acknowledgements

This work was supported by the grants from the National Institutes of Health Common Fund 4D Nucleome Program (U01CA200059) and National Institutes of Mental Health (U01MH106883) to PJP.

