## Supplemental figures for "HiTea: a computational pipeline to identify non-reference transposable element insertions in Hi-C data"

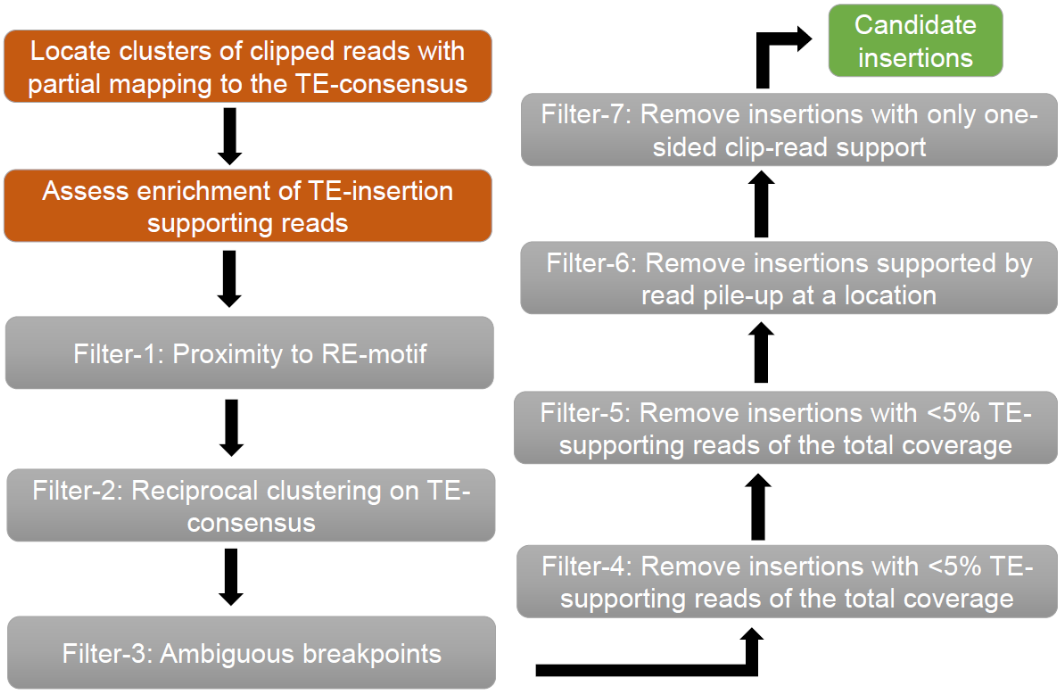


**Supplementary Figure 1: HiTea workflow and filters.** HiTea starts by first identifying clipped-reads with partial mapping to the TE-consensus assembly. All reads where the clipped-coordinates fall within 2bp are grouped together as referred as clusters of reads. Clusters supported by at least 2 reads are assessed for enrichment of all the TE-supporting reads (clipped reads + RAM pairs) at the locus (steps are highlighted in brown). Next, HiTea uses series of filtering steps to remove potential false positive instances (highlighted in gray). The filtering steps take into account the presence of restriction endonuclease (RE) site at the breakpoint, reciprocal clustering on TE-consensus and so on. When reads pile-up at the same location, i.e. with same start, clip and read-orientation, such clusters are omitted as potential amplification related artifacts. Finally, when clip-read support is present on either left or right-hand site, the clusters are filtered out as low confidence instances

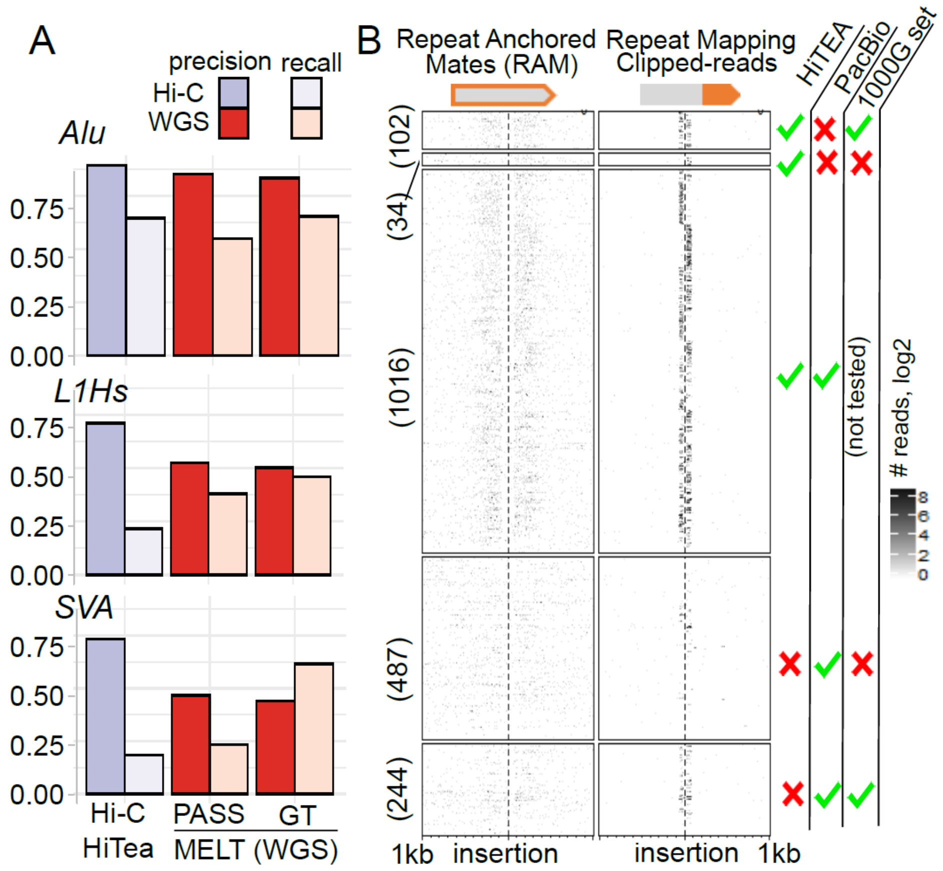


**Supplementary Figure 2: HiTEA performance at 600M sequencing depth (vs 1.4B in Figure 2). (A)** Precision and recall for detecting insertions of Alu, L1Hs and SVA families using HiTea (on Hi-C) and MELT (on WGS) for the samples with 600M RPs. PASS and GT refer to the more and less stringent call sets, respectively, in MELT. **(B)** 5' end (read start) coverage for the RAMs whose mates map to the TE consensus (left) or reads whose clipped-sequences map to the TE consensus (right). The insertions are grouped according to the criteria shown on the right. PacBio is the reference set constructed using PacBio HiFi reads; 1000G set refers to insertions detected in the 1000 Genome data by MELT

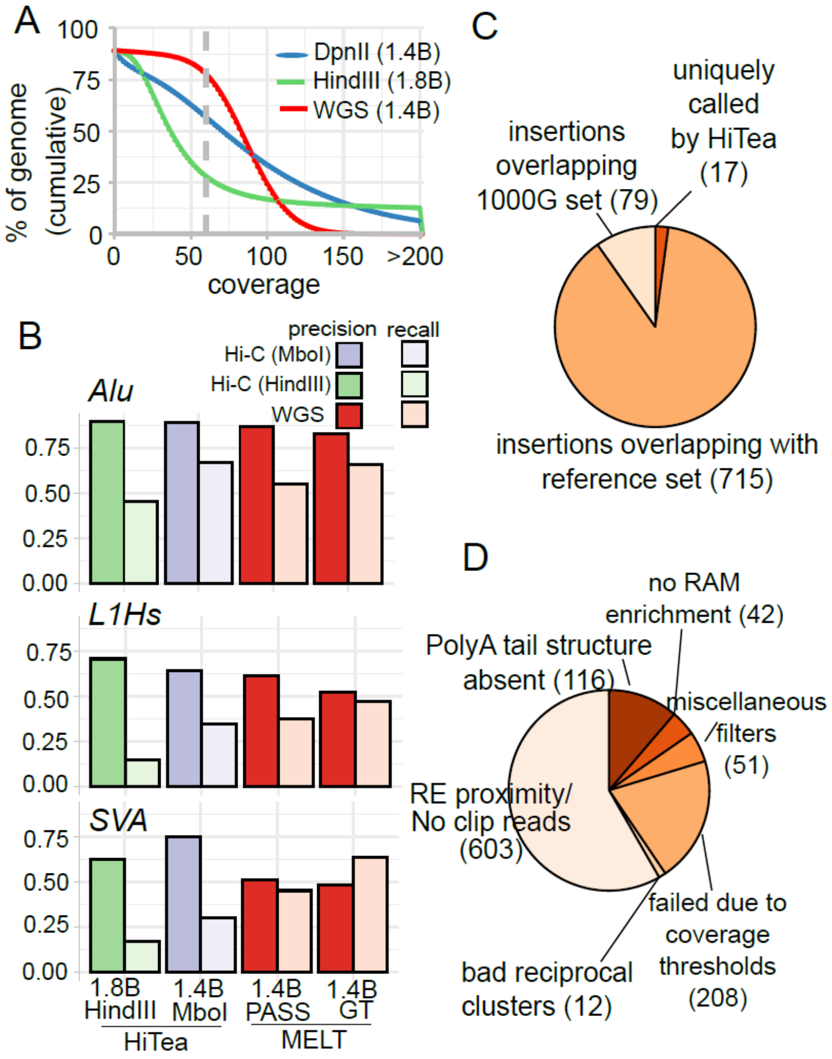


**Supplementary Figure 3: Performance comparison between HindII- and MboI-digested Hi-C data. (A)** Coverage comparison for Hi-C samples derived after HindIII or MboI digestion as well as for WGS. Gray line represents 60X coverage. **(B)** Precision and recall comparison in identifying TE insertions. **(C)** Overlap of TE insertions determined by HiTea (on HindIII digested data) with reference sets. **(D)** Filters that led to non-identification of PacBio-reference insertions in the HindIII digested Hi-C data.

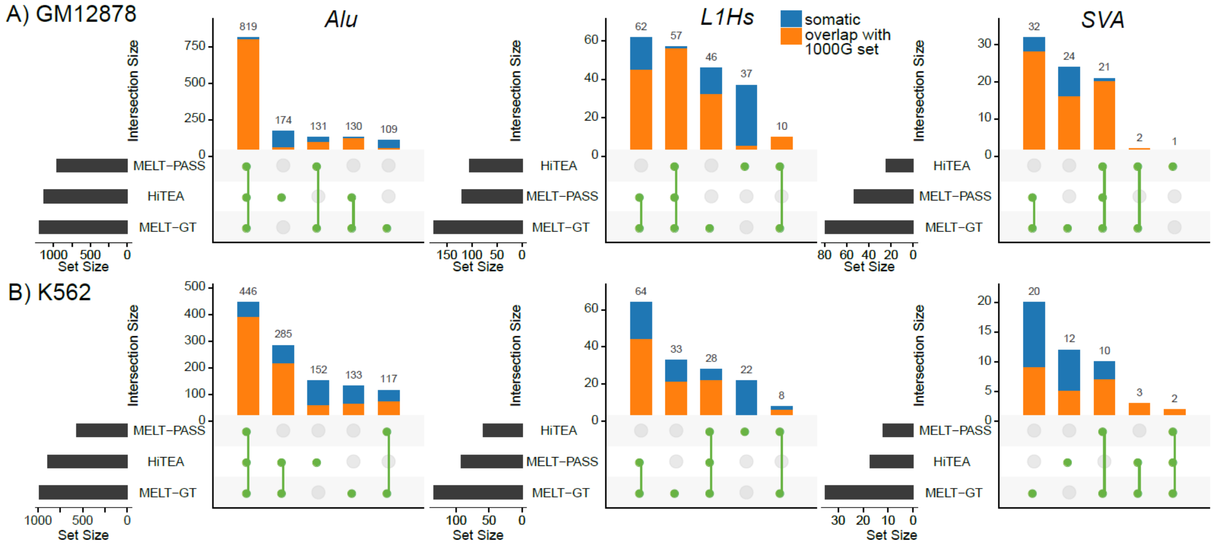


**Supplementary Figure 4: Performance comparison between HiTea and MELT. (A)** Intersection of insertion candidates identified by HiTea (on MboI digested data) and MELT (on WGS data) at 1.4B sequencing depth for GM12878 cell line. Insertions overlapping with 1000G set are displayed in orange. **(B)** Similarly, for K562 (1.2B)

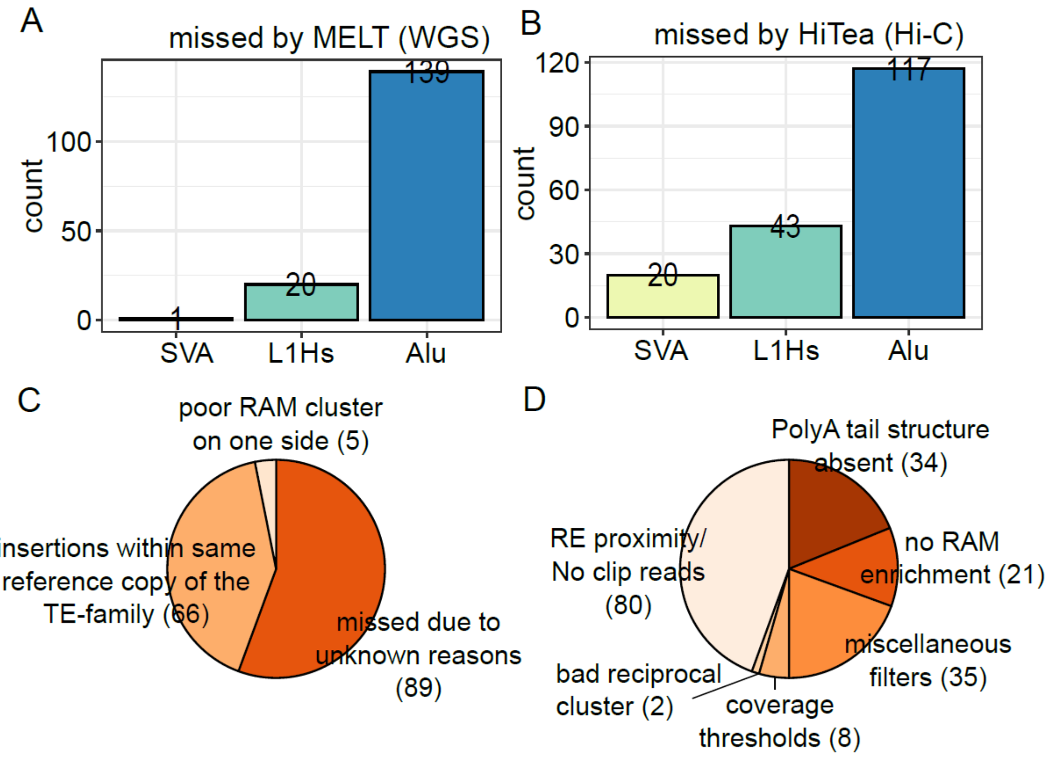


**Supplementary Figure 5: PacBio reference insertions missed by MELT or HiTea. (A, B)** TE insertions missed by one algorithm but missed by the other. **(C, D)** Features that likely led to the non-identification of the insertions by either MELT or HiTea, respectively. As the filtering steps in MELT are not disclosed, the missed insertions are grouped based on orthogonal features observed from the data in (C).
